# Supporting per-locus substitution rates improves the accuracy of species networks and avoids spurious reticulations

**DOI:** 10.1101/2022.01.16.476511

**Authors:** Zhen Cao, Huw A. Ogilvie, Luay Nakhleh

**Affiliations:** Department of Computer Science, Rice University, Texas, USA

**Keywords:** Phylogenetic networks, multispecies coalescent, phylogenomics, rate heterogeneity

## Abstract

The development of statistical methods to infer species phylogenies with reticulation (species networks) has led to many discoveries of gene flow between distinct species. However, because the dimensionality of species networks is not fixed, these methods may compensate for kinds of model misspecification, such as assuming a single substitution rate for all genomic loci, by increasing the number of dimensions beyond the true value. The popular full Bayesian species network method `MCMC_SEQ` has previously made this assumption, so we have added support for the proven Dirichlet model for per-locus rates to enhance its accuracy and avoid spurious results. We studied the effects of this model using simulation and an empirical dataset from *Heliconius* butterflies.

We found that assuming a single substitution rate applies to all loci leads to the inference of spurious reticulation in simulated and empirical datasets when a full Bayesian method is used, however, the summary method `InferNetwork_ML` is robust to per-locus variation in substitution rates when set to ignore gene tree branch lengths. Our implementation of the model resolves this misspecification and successfully converges to the true species networks. It also infers far more accurate gene trees than assuming a single rate, or independent inference of gene trees.

Our implementation of the Dirichlet per-locus rates model is now available in PhyloNet, a software package for phylogenetic inference, open source on GitHub https://github.com/NakhlehLab/PhyloNet.

## Introduction

Our understanding of evolutionary history is limited by the models used to represent those histories. The most common model is a phylogenetic *tree*, which may be used to model among other things the histories of genes (by which we mean discrete parts of a genome called loci) and species. When a phylogenetic tree is used to model the history of a set of genes related by common descent it is known as a gene tree. When used to model species histories and hence known as species trees, branching events represent instances of speciation, or the splitting of one ancestral species into descendant species. Species trees by themselves do not permit reticulation, instead building in an assumption that once a species is divided, no gene flow (where individuals from different contemporaneous species may produce offspring) between the divided populations will occur subsequently. Forms of gene flow that may occur are introgression (the transmission of genes from one species to another) and hybrid speciation (where the offspring of parents from different species form a new species). As a result, the discovery of many instances of reticulate evolution was delayed until the first methods were developed to detect gene flow.

The development of methods to infer reticulation evolution among eukaryotic nuclear genomes is particularly challenging because incomplete lineage sorting (ILS) causes incongruence of gene and species phylogenies even in the complete absence of reticulation [15]. Gene duplication and loss may also result in incongruence without reticulation [19]. Nevertheless, methods which discriminate between these processes have been developed—for example, the ABBA-BABA statistical test enabled the detection of introgression from Neanderthals into modern humans [8].

The identification of introgression in eukaryotes motivated the development of combinatorial and statistical methods that use phylogenetic *networks* to model the evolutionary history of species (henceforth *species networks*) [25, 24, 23, 22, 28, 27, 26, 6]. Such methods generalize the multispecies coalescent (MSC) model, which specifies a probability distribution over gene trees given a species tree and a demographic model. The generalized model instead uses a species network and is known as the multispecies network coalescent (MSNC) model. MSNC methods enabled the discovery of more instances of introgression [12, 7], demonstrating the importance of tractable and reliable methods for detecting and characterizing reticulate evolution.

One class of MSC and MNSC methods is multilocus methods, which use alignments of genes as input. These methods assume recombination is frequent between genes and limited within them, so that a single gene tree can be inferred for individual alignments, and the histories of different alignments are independent samples from the underlying distribution of gene trees. These methods may be further divided into summary methods and full methods. Summary methods use previously inferred gene trees to estimate the species phylogeny, whereas full methods jointly estimate the species phylogeny and gene trees in a single step.

The rate at which genes evolve is known as the substitution rate and until now the effects of substitution rate heterogeneity on network inference methods have not been explored. Substitution rates are suspected to vary greatly between loci [3], changing the branch lengths of gene trees when measured in the number or expected number of substitutions. Summary methods, such as `InferNetwork_ML` [24], are usually set to ignore branch lengths, and so should be robust to differences in true branch lengths, although perhaps not to the increased error which is inversely correlated with branch length. However, the full method `MCMC_SEQ` [22] previously assumed all loci evolve at the same substitution rate and may be misled since the species network probability is dependent on coalescent times within each gene tree which are scaled by the substitution rate at each locus. This is a particular concern when reticulations are permitted by the model and method, since spurious reticulations may be added to the species network in order to account for the apparently (but not actually) different coalescent times.

Herein we conduct a simulation study using the above methods to understand their behaviour and susceptibility to error in the presence of substitution rate variation. We have also enhanced `MCMC_SEQ` by incorporating support for varying substitution rates with a fixed mean and an implied flat Dirichlet prior by adding a delta exchange operator (DEO) to that method. Throughout the rest of the paper, in the context of simulation and inference, we will refer to this as the “Dirichlet rates” (DR) model, as opposed to a single rate (SR) for all loci. Both operator and model of substitution rate variation were originally introduced in BEAST [2]. We show that allowing for substitution rate variation dramatically improves the accuracy of species network inference when rate variation is present in the data, and does not meaningfully reduce the accuracy of inference even when all loci have the same rate.

## Materials and methods

### Definitions

We use the same definitions as in the original publication of `MCMC_SEQ` [22], with necessary modifications to implement the DR model using the DEO operator.

### Phylogenetic networks and their parameters

A *phylogenetic network* Ψ on set 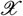 of taxa, is a rooted directed, acyclic graph (DAG) with vertices *V* (Ψ) = {*s, r*}∪*V_L_* ∪*V_T_* ∪*V_N_*, where

- *indeg*(*s*) = 0 and *outdeg*(*s*) = 1 (*s* is a special node, that is the parent of the root node, *r*);
- *indeg*(*r*) = 1 and *outdeg*(*r*) = 2 (*r* is the *root* of Ψ);
- ∀*v* ∈ *V_L_*, *indeg*(*v*) = 1 and *outdeg*(*v*) = 0 (*V_L_* are the *external tree nodes, or leaves*, of Ψ);
- ∀*v* ∈ *V_T_*, *indeg*(*v*) = 1 and *outdeg*(*v*) ≥ 2 (*V_T_* are the *internal tree nodes of* Ψ); and,
- ∀*v* ∈ *V_N_*, *indeg*(*v*) = 2 and *outdeg*(*v*) = 1 (*V_N_* are the reticulation nodes of Ψ).

The set of edges, *E*(Ψ) ⊆ *V* × *V*, consists of *reticulation edges*, whose heads are reticulation nodes, *tree edges*, whose heads are tree nodes, and special edge (*s, r*) ∈ *E*. Besides, the *leaf-labeling* function, which is a bijection from *V_L_* to 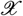, is 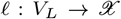. Each node in *V* (Ψ) is associated with a divergence time, and each edge in *E*(Ψ) is associated with a population size. The edge length of *er*(Ψ) = (*s, r*) is infinite so that all lineages entering it coalesce on it eventually. Each pair of reticulation edges *e*_1_ and *e*_2_ that share the same reticulation node, is associated with an inheritance probability, *γ*, such that 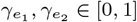 with 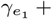 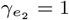. We denote by Γ the vector of inheritance probabilities corresponding to all the reticulation nodes in the phylogenetic network (for each reticulation node, Γ has only one of the values of the two incoming edges).

Given a phylogenetic network Ψ representing the topology, species divergence times and population size parameters, we use the following notation:

- Ψ_*top*_: The leaf-labeled topology of Ψ; that is, the pair (*V, E*) along with the leaf-labeling *ℓ*.
- Ψ_*ret*_: The number of reticulation nodes in Ψ. Ψ*_ret_* = 0 when Ψ is a phylogenetic tree.
- Ψ*_τ_* : The species divergence time parameters of Ψ. Ψ*_τ_* ∈ (ℝ^+^)^|*V* (Ψ)|^.
- Ψ_*θ*_: The population size parameters of Ψ. Ψ*_θ_* ∈ (ℝ^+^)^|*E*(Ψ)|^. For each branch, the corresponding *θ* equals 4*N_e_μ*, where *N_e_* is the effective population size for that branch, and *μ* is the mutation rate per site per generation.

### Bayesian Formulation and Inference

The data in our case is a set 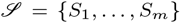 where *S_i_* is a DNA sequence alignment from locus *i*. A major assumption is that there is no recombination within any of the *m* loci, yet there is free recombination between loci. The model 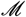 consists of a phylogenetic network Ψ (the topology, divergence times, and population sizes) and a vector of inheritance probabilities Γ. The posterior distribution of the model is given by

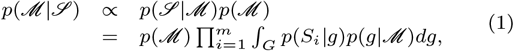

where the integration is taken over all possible gene trees. The term *p*(*S_i_*|*g*) gives the gene tree likelihood, which is computed using Felsenstein’s algorithm [5] assuming a model of sequence evolution which includes a substitution rate *μ*, and 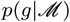 is the probability density function for the gene trees, which was derived for the cases of species tree and species network in [18] and [24], respectively.

The integration in Eq. (1) is computationally infeasible except for very small data sets. Furthermore, in many analyses, the gene trees for the individual loci are themselves a quantity of interest. Therefore, to obtain gene trees, we sample from the posterior distribution as given by

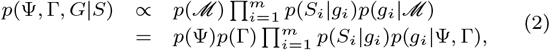

where *G* = (*g*_1_*,..., g_m_*) is a vector of gene trees, one for each of the *m* loci. This co-estimation approach is adopted by the Bayesian methods *BEAST [9] and BEST [11], both of which co-estimate species and gene trees, and `MCMC_SEQ` [22], which co-estimates phylogenetic networks and gene trees.

### Implementing the Dirichlet rates model

We adapted a proposal, delta exchange operator (DEO), from BEAST [2] to sample substitution rate of each locus in the code base of `MCMC_SEQ` [22] in PhyloNet.

Instead of assuming a common substitution rate for all loci, we now incorporate a vector of substitution rates *M* = *μ*_1_*, μ*_2_*,.., μ_m_* in our model. Using a multiplier *δ* and a vector of weights for each substitution rates *W* = *w*_1_, *w*_2_,.., *w_m_*, the operator proposes changing two rates in *M* at a time. The first step is to select two indices of substitution rates *d*_1_, *d*_2_ with weights 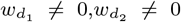. Then, the operator computes a value *x* = *r* × *δ*, where *r* is a random number. Finally, we have 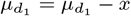, and 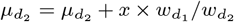 as proposed values for substitution rates. Note that this operator holds the sum/weighted sum of substitution rates *μ*_1_*, μ*_2_*,.., μ_m_* constant. An example is in Fig. 1.

**Fig. 1:**
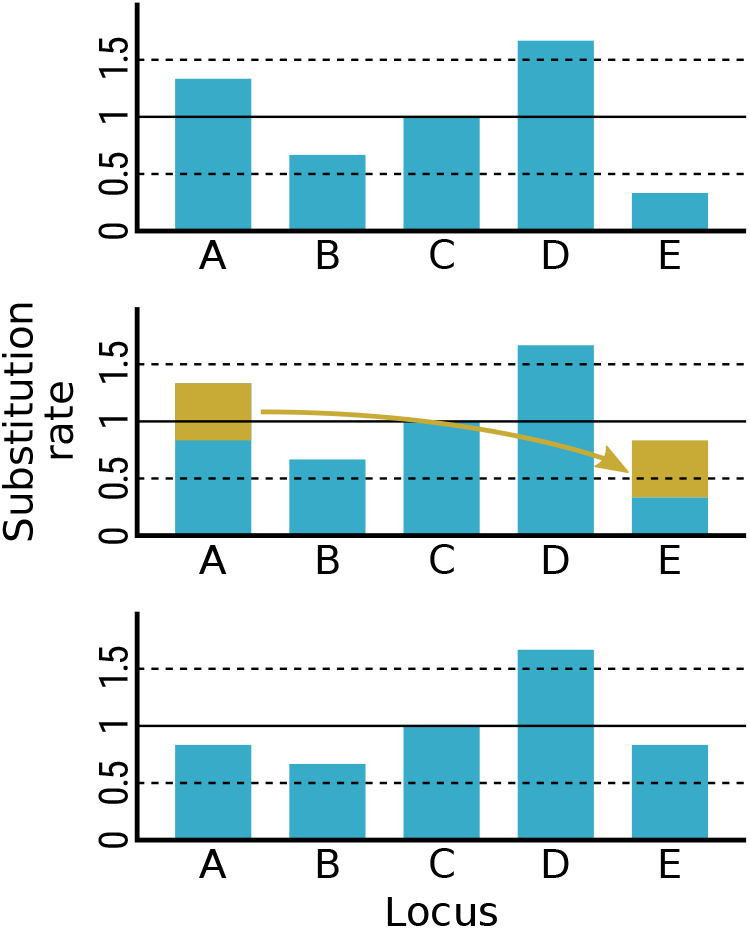
Example proposal by the Delta Exchange Operator (DEO). The current state has a mean substitution rate of 1 (solid horizontal line, top). The substitution rate of locus A is proposed to be decreased and the rate of locus E is proposed to be increased by an equal amount (amounts and transfer shown in gold, middle). If accepted, this proposal maintains the mean substitution rate of 1 (solid horizontal line, bottom).

### Simulating phylogenies and sequence alignments

For each replicate, we used the program `ms` [10] to simulate 100 conditionally independent gene trees for each exemplar species phylogeny (Fig. 2). The number of replicates was 10 for each phylogeny, and the following commands were used for each replicate respectively.

**Fig. 2:**
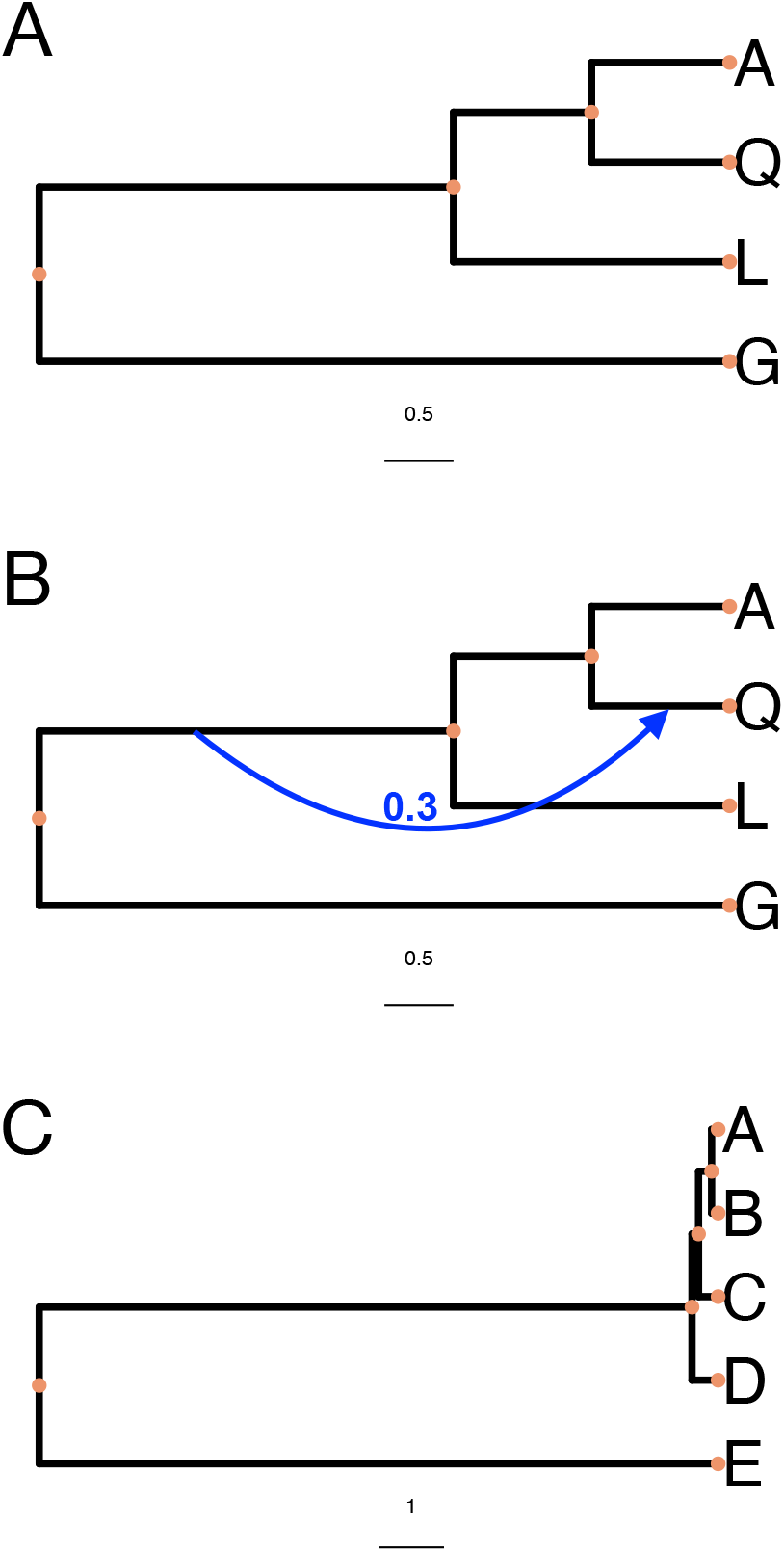
Exemplar phylogenies used in our simulation study. These were chosen from a displayed subtree of anophiline mosquitos (A), a displayed subnetwork of the same taxa (B) and a species tree in the “anomaly zone” (C). Taxon labels A, Q, L and G refer to *Anopheles arabiensis*, *A. quadriannulatus*, *A. melas* and *A. gambiae* respectively. The blue edge highlights which branch leading to the reticulation node in (B) has the smaller inheritance probability *γ* = 0.3. Branch lengths are in coalescent units.

~~~
ms 3 100 -T -I 3 1 1 1 -ej 0.75 3 2 -ej
1.0 2 1
ms 4 100 -T -I 4 1 1 1 1 -ej 0.5 2 1 -ej
1.0 3 1 -ej 2.5 4 1
ms 5 100 -T -I 5 1 1 1 1 1 -ej 0.05 5 4
-ej 0.15 4 3 -ej 0.2 3 2 -ej 5.2 2 1
~~~

We generated two data sets for each replicate for comparison, one with homogeneous rates (the SR model), the other with heterogeneous rates (the DR model). At each locus, we used program `seq-gen` [17] to generate 500bp sequence alignments, under a GTR model with base frequencies 0.2112, 0.2888, 0.2896, 0.2104 (A, C, G, T) and transition probabilities 0.2173, 0.9798, 0.2575, 0.1038, 1.0, 0.207 (A to C, A to G, A to T, C to G, C to T, T to G). We specified the population mutation rate to be *θ* = 0.036, and therefore the simulated gene trees were scaled by 0.018 to convert their branch lengths into expected substitutions per site. The command below was used to generate homogeneous sequence data with substitution rate 1 for all loci.

~~~
seq-gen -mGTR -s0.018 -f0.2112,0.2888,
0.2896,0.2104 -r0.2173,0.9798,0.2575,
0.1038,1.0,0.207 -l500
~~~

To generate data with heterogeneous rate per locus, we further sampled a vector of scales *s*_1_, *s*_2_,..., *s_m_* under the Dirichlet distribution *f* (*s*_1_, *s*_2_,..., *s_m_*, 1, 1,..., 1), where the concentration parameter is a vector of *m* values all set to 1. Then we scaled gene trees by 0.018 × *s_i_* as part of the `seq-gen` command to generate sequences. As with the homogenous rates data, sequence alignments of 500bp length were simulated.

### Species phylogeny and gene tree inference

We estimated the gene tree phylogeny for each locus independently of the species tree using IQ-TREE version 1.6.10 [16], and rooted at outgroup. Then we used `InferNetwork_ML` to infer species phylogeny with the number of runs set to 50 and the number of networks returned set to 10, with branch length and inheritance probabilities post-optimized. We set the maximum number of reticulations to 0, 1, 2 or 3 to calculate the maximum likelihood topology and log-likelihood for the network with the corresponding number of reticulations. We ran `MCMC_SEQ` on multi-locus alignment data with chain lengths set to 50,000,000, the burn-in set to 5,000,000 and the sample frequency set to 1 in 5,000.

### Extracting *Heliconius* sequence alignments

We utilized whole genome alignments of *Heliconius* sequences [4]. We extracted 10kb windows spaced at 50kb intervals using *makewindows* in *bedtools* v2.29.2. Then, we used *hal2maf* v2.1 to obtain *MAF* alignments based on reference genome. We converted alignments to fasta format with tool *msa_view*. We then filtered loci with maximum Jukes Cantor distance between species greater than 0.2, restricted the taxa to 2 sets of species, including set 1 (*H.erato.demophoon*, *H.hecalesia*, *H.melpomene*), and set 2 (*H.melpomene*, *H.timareta*, *H.numata*), and an outgroup *Agraulis vanillae*, and filtered loci with missing species. Finally, we selected 100 loci at random and truncated each loci to 500 bps.

We inferred a gene tree for each locus with IQ-TREE, and rooted them using the outgroup taxon *Agraulis vanillae*. We ran `InferNetwork_ML` on estimated gene trees with the number of runs 50 and number of networks returned 10 with branch lengths and inheritance probabilities post optimized, and set the maximum number of reticulations to 0-3 to decide the number of reticulations by observing the amount of increment in log likelihood.

We ran 10 chains of `MCMC_SEQ` with 80,000,000 iterations, 8,000,000 burn-in, 5,000 sample frequency, allowing one population mutation rate per branch for both models of uniform substitution rates and variable substitution rates. Each chain was started with a unique random seed to ensure samples from the different chains were independent. We summarized the species network as the topology with the highest marginal probability from the samples across all chains after burn-in, and summarized continuous parameters using their mean and standard deviation.

### Quantitative measures of accuracy and error

Normalized Robinson-Foulds distance [20] of trees *T*, 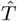 is defined as the number of clades present only in one of *T*, 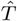 normalized by the maximum number of different clades, where *T* is the true tree, 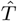 is the estimated tree.

Normalized rooted branch score [9] is defined as 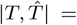 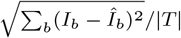, where *I_b_* is the branch length of the true tree, 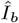 is the branch length of the estimated tree, for a branch *b* in the set of all branches in the tree, and |*T*| is the total length of the true tree.

The relative error of a substitution rate is 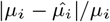, and the average relative error is 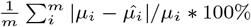.

## Results

### Simulation study

Using simulation, we studied the ability of the summary maximum likelihood method `InferNetwork_ML` and the full Bayesian method `MCMC_SEQ` to reconstruct three exemplar species phylogenies from gene trees and sequence alignments simulated based on those phylogenies either under the DR or SR models. The basis of first exemplar phylogeny is a tree displayed by an inferred species network of anopheline mosquitoes pruned to the taxa *Anopheles arabiensis*, *A. quadriannulatus*, *A. melas* and *A. gambiae* [Fig. 2A, 7]. The basis of the second exemplar phylogeny is a displayed network of the same pruned species network (Fig. 2B), and the third is a previously published species tree in the anomaly zone [Fig. 2C, 13]. We assessed how the accuracy of species network topologies was impacted by rate heterogeneity using the network topology distance metric [14] (henceforth referred to as network distance) which is more fine-grained than simple identity. To assess the accuracy of gene trees we applied the normalized versions of the commonly used Robinson-Foulds and branch score measures (henceforth referred to as nrRF and nrBS respectively).

### Summary methods are robust to substitution rate heterogeneity

Maximum likelihood methods of species network inference, unlike Bayesian methods, are not inherently able to estimate the number of reticulations present. This is because a network of higher likelihood should always be possible to find by increasing the number of reticulations *k*, even beyond the true *k* [1]. Therefore a maximum likelihood search should return a network where *k* is equal to the maximum permissible value *k_max_* set by the user or implementation. When *k_max_* is set equal to the true *k*, introducing rate heterogeneity has only a very minor negative effect on species network accuracy when gene trees are estimated (Fig. 3), despite a more substantial decrease in the accuracy of inferred gene tree topologies (Fig. 4). When the true phylogeny is a network, using estimated gene trees has a very negative effect (scenario 2 in Fig. 3), but adding rate heterogeneity does not make it any worse.

**Fig. 3:**
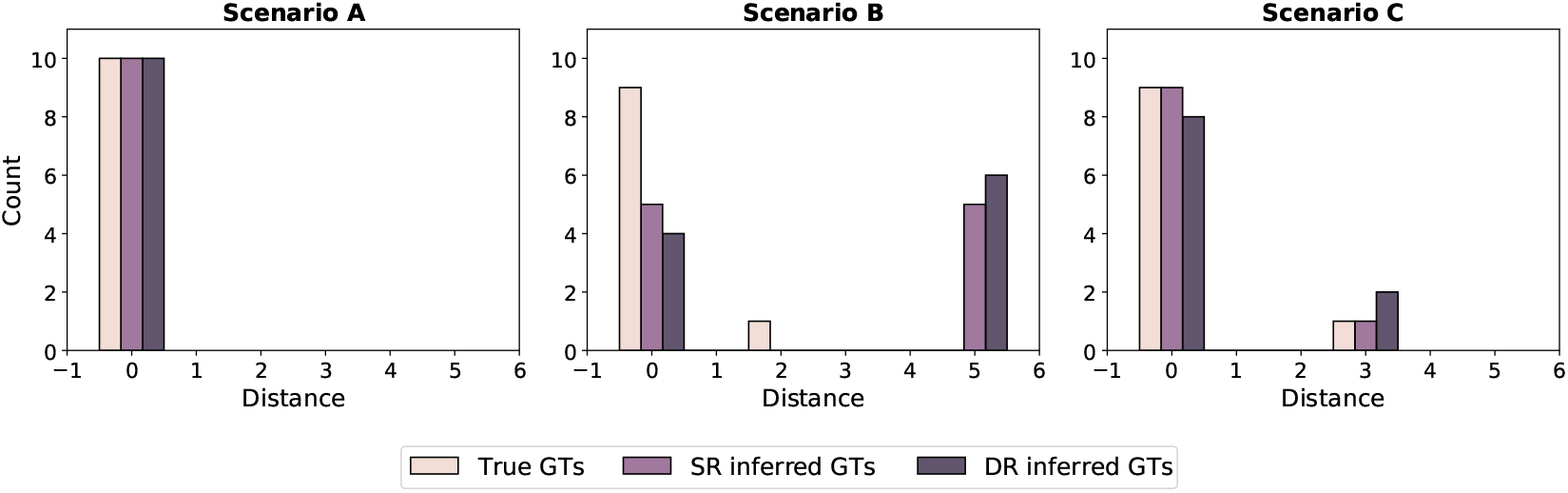
The effect of per-locus substitution rate variation on the accuracy of species phylogenies inferred by the maximum likelihood summary method `InferNetworks_ML`. Each bar represents the number of replicates a given network distance from the true species network, for the three exemplar species phylogenies (scenarios) A, B and C. Gene trees used as input were either the true gene trees (True GTs), or inferred using IQ-TREE from sequence data simulated along those trees under a single rate model (SR inferred GTs), or a Dirichlet rates model (DR inferred GTs).

**Fig. 4:**
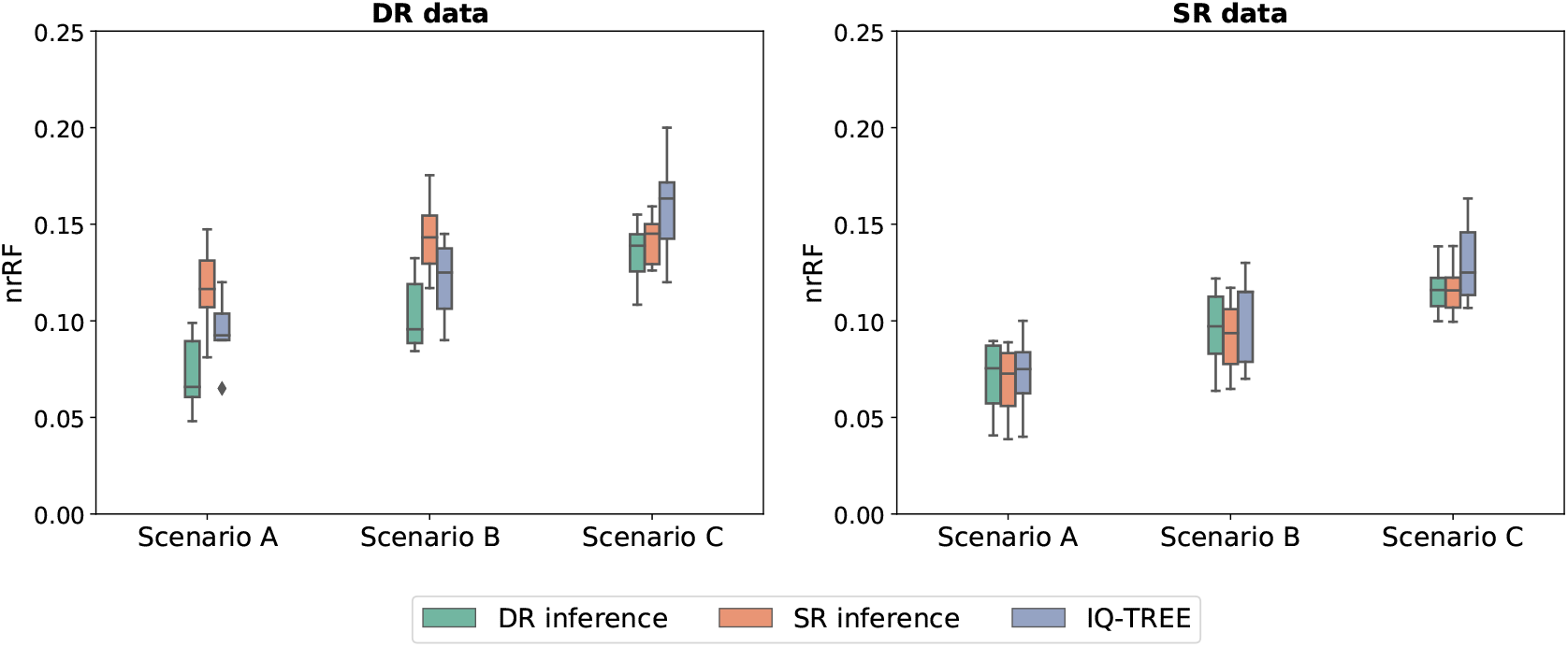
The effect of per-locus substitution rate variation on the accuracy of gene tree topologies inferred independently or using a full Bayesian approach. IQ-TREE was used to infer gene trees independently of each other or the species phylogeny. `MCMC_SEQ` was used to infer gene trees together with the species phylogeny from multiple sequence alignments in a full Bayesian approach, under either single rate (SR) or Dirichlet rates (DR) models of per-locus substitution rate variation. Sequence alignment data were also simulated under either SR or DR models.

The noticeable difference between the accuracy of species networks and trees inferred from estimated gene trees cannot be attributed to differences in gene tree error, as the nrRF of IQ-TREE is 0.122 for the species network, and is 0.093 and 0.159 for the anophiline and anomaly zone species trees respectively under rate heterogeneity (Fig. 4). Adding rate heterogeneity to the true gene trees has no effect since this summary method ignores branch lengths and the true gene tree topologies are unaffected by rate heterogeneity.

The original publication on maximum likelihood species network inference [24] suggested increasing *k* until the increase in likelihood becomes negligible. This creates the appearance of a “shoulder” when the maximum likelihood is drawn as a function of *k*. The number of reticulations is able to be identified when the phylogeny is not in anomaly zone, since we observe little growth in the likelihood after the point of the true number of reticulations (Fig. 5 scenario 1 and 2). While it is concerning that there is steady increase of log likelihood in anomaly zone (Fig. 5 scenario 3), and we leave this to future work.

**Fig. 5:**
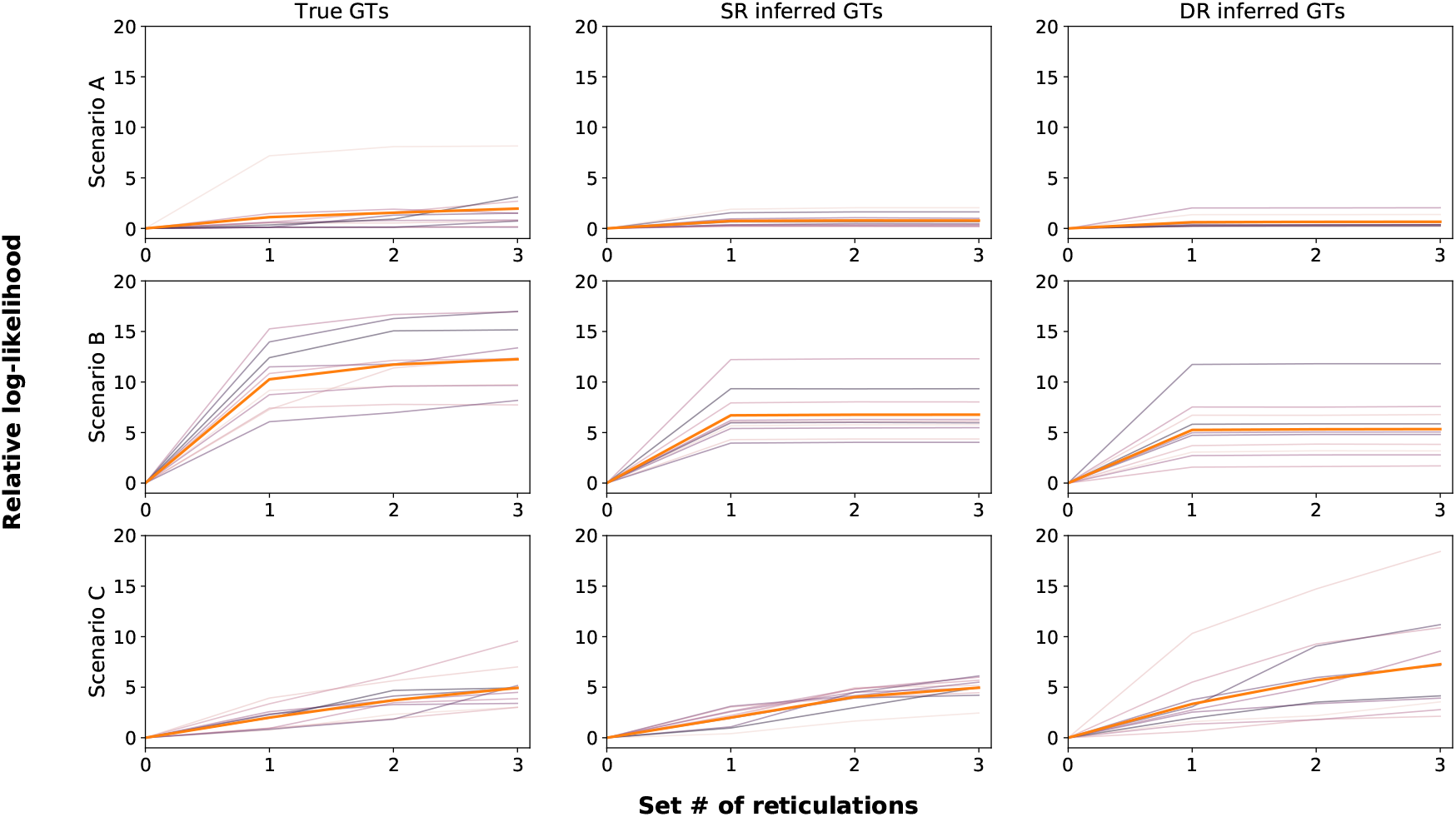
Log-likelihood increase of species networks identified using the maximum likelihood summary method `InferNetworks_ML`. The log-likelihood for a given maximum number of reticulations is relative to the log-likelihood of the zero reticulation maximum likelihood network (i.e. the species tree). Gene trees used as input were either the true gene trees (True GTs), or inferred using IQ-TREE from sequence data simulated along those trees under a single rate model (SR inferred GTs), or a Dirichlet rates model (DR inferred GTs). Thick orange lines show the average increase relative to zero reticulations, all other lines show the increase for each individual replicate.

### Allowing for differing substitution rates is needed for full Bayesian methods

Unlike maximum likelihood methods, the number of reticulations is identifiable when using Bayesian methods. So for the full Bayesian method `MCMC_SEQ` we evaluated the accuracy of the *maximum a posteriori* (MAP) species network without pre-specifying the number of reticulations. When data was simulated under the SR model and lacked rate heterogeneity, using the DEO implementation of the DR model for inference did not have a substantial negative effect on topological accuracy. However, when data was simulated under the DR model, and the DEO implementation was not used and so the SR model was used for inference, we observed a large decrease in accuracy (Fig. 6 top row). Further investigation revealed that the basis for this error was the inference of reticulations beyond the true number of reticulations (Fig. 6 bottom row). Similar results were observed for the species network with the highest probability marginalized over branch lengths and other continuous parameters although with lower accuracy when the DEO implementation was used, likely because this point estimate of the network requires greater time to converge (Fig. S1).

**Fig. 6:**
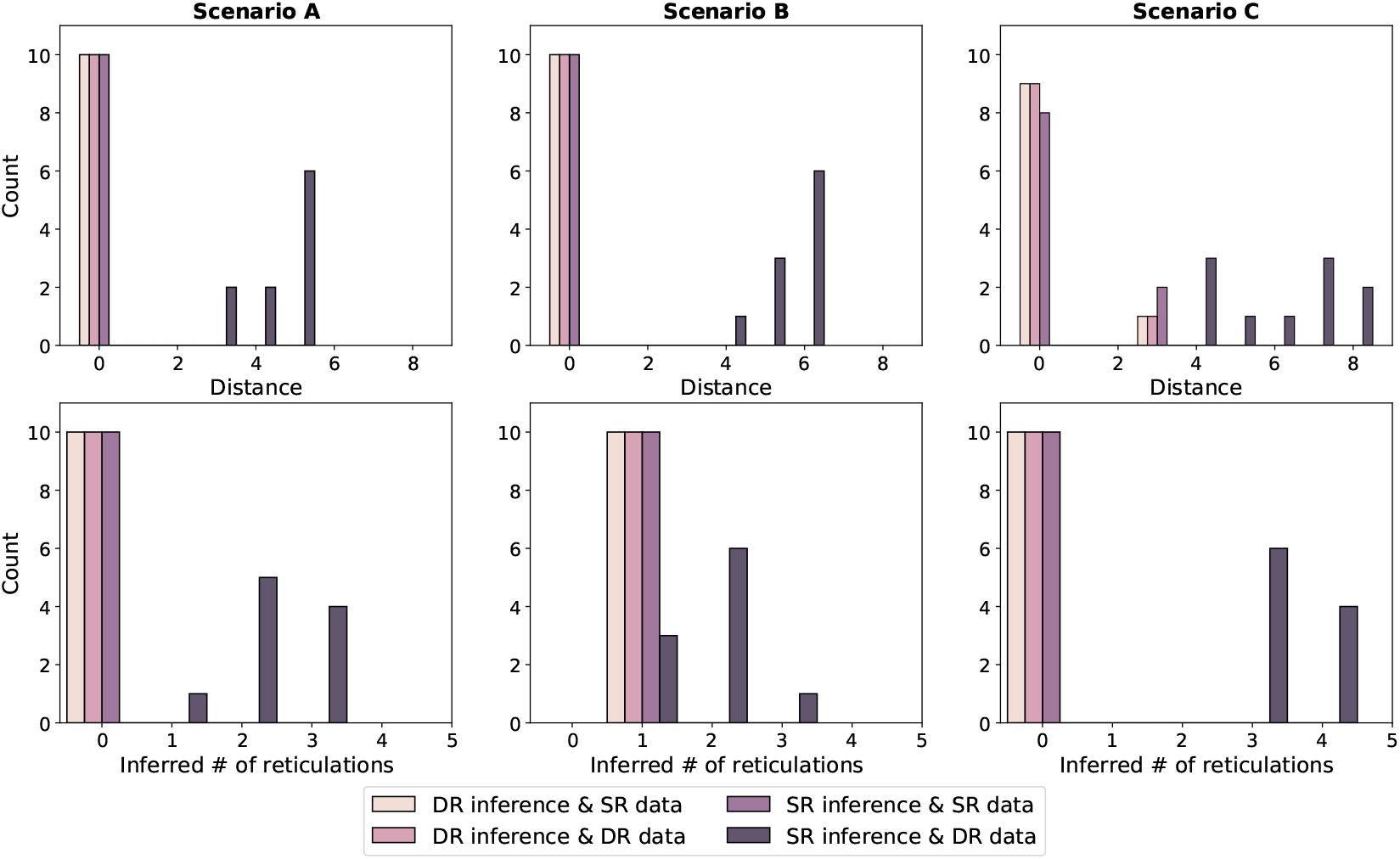
The effect of per-locus substitution rate variation on the accuracy of species phylogenies inferred using a full Bayesian approach. `MCMC_SEQ` was used to infer the species phylogeny together with the gene trees from multiple sequence alignments in a full Bayesian approach, under either single rate (SR) or Dirichlet rates (DR) models of per-locus substitution rate variation. Sequence alignments were also simulated under either SR or DR models. Each bar represents the number of replicates a given network distance from the true species network, for the three exemplar species phylogenies (scenarios) A, B and C.

### Full Bayesian methods always outperform independent gene tree estimation

Full Bayesian estimation of the species phylogeny and gene trees improves gene tree accuracy compared with inference of gene trees independently prior to species tree inference [21]. Methods like `MCMC_SEQ` may therefore be used to infer more accurate gene trees than `IQ-TREE`, the program we used for the summary method workflow. When data was simulated without rate heterogeneity under the SR model, `MCMC_SEQ` performed similarly to `IQ-TREE` when inferring gene tree topologies regardless of whether the DEO implementation of the DR model was enabled. However, when data was simulated under the DR model with rate heterogeneity, using the DR model for inference with `MCMC_SEQ` consistently outperformed the SR model or `IQ-TREE` (Fig. 4).

When branch lengths are considered in addition to topology using the nrBS measure, the difference between the methods is dramatic. Regardless of whether data was simulated under the DR or SR models, and whether the DEO implementation of the DR model was enabled, the full Bayesian `MCMC_SEQ` method massively outperformed independent inference of gene trees using `IQ-TREE`. Although full Bayesian methods were always superior to independent inference, when data was simulated under the DR model, using the DEO implementation of the DR model for inference massively outperformed the SR model as well (Fig. 7).

**Fig. 7:**
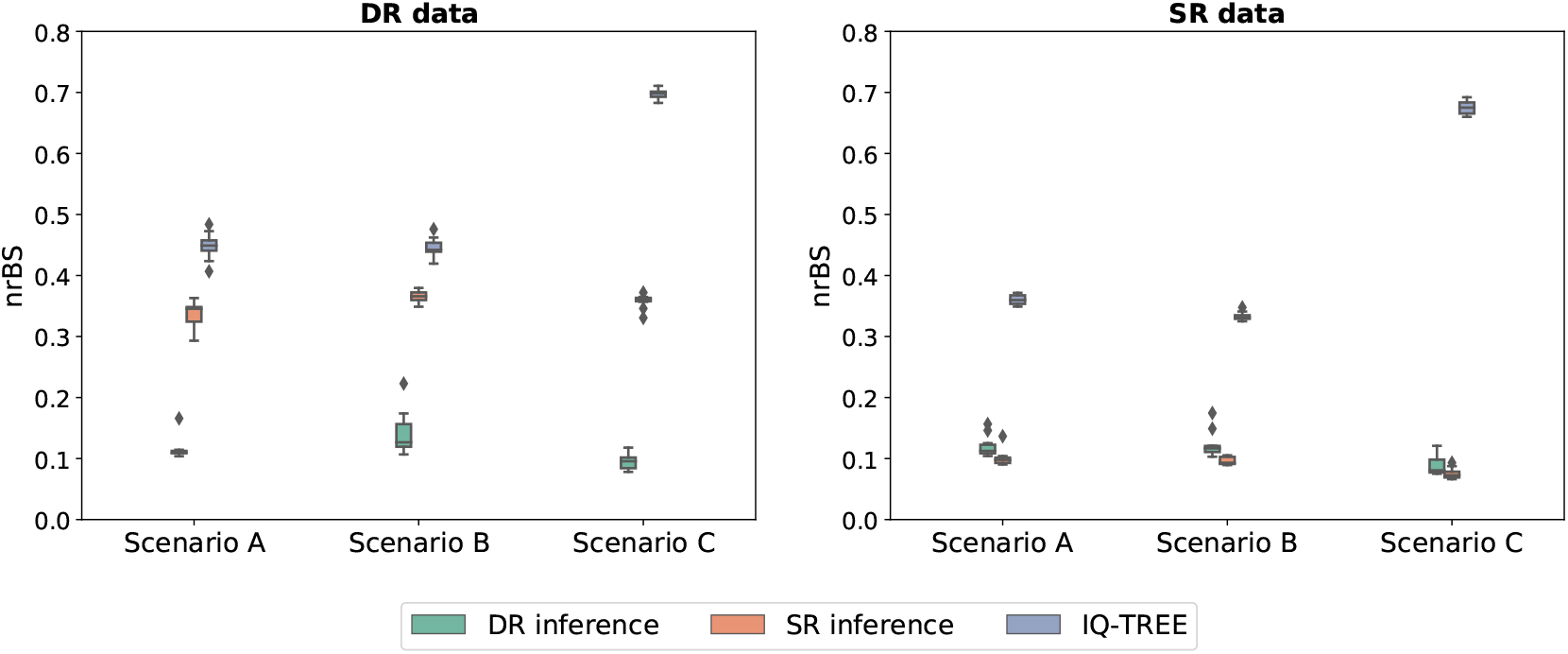
The effect of per-locus substitution rate variation on the accuracy of gene tree branch lengths inferred independently or using a full Bayesian approach. `IQ-TREE` was used to infer gene trees independently of each other or the species phylogeny. `MCMC_SEQ` was used to infer gene trees together with the species phylogeny from multiple sequence alignments in a full Bayesian approach, under either single rate (SR) or Dirichlet rates (DR) models of per-locus substitution rate variation. Sequence alignment data was also simulated under either SR or DR models.

The relative substitution rates estimated under the DR model using the DEO implementation in `MCMC_SEQ` were strongly correlated with the true simulated rate (Fig. 8), although the average relative error was 35.98%, reflecting the limited information present in each individual sequence alignment. Estimation of relative rates is not possible using `IQ-TREE` due to the confounding of rates and time.

**Fig. 8:**
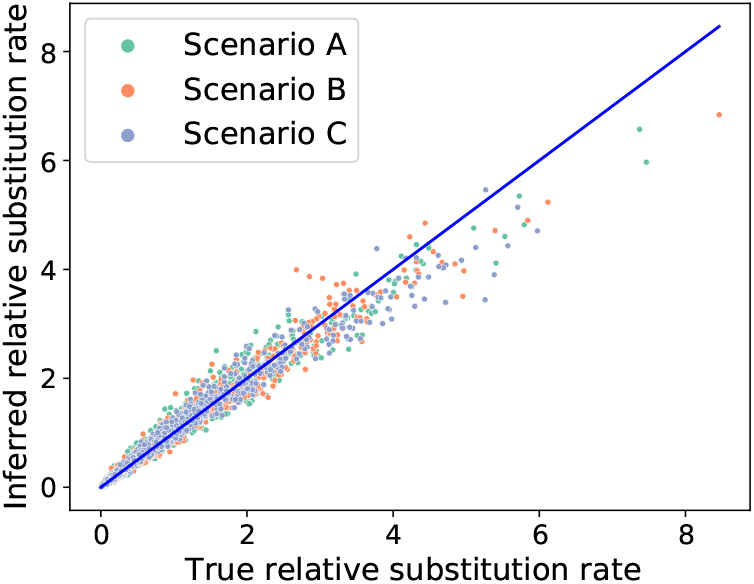
Per-locus substitution rate estimates using the Dirichlet rates model of per-locus substitution rate variation. Our implementation of this model in `MCMC_SEQ` was used to estimate the rates from sequence alignments simulated using the same model, from each of the three exemplar phylogenies (scenarios A, B and C). Rates along the diagonal exactly match the true rates used for simulation.

### When not accounting for rate heterogeneity, *Heliconius* phylogenies may be erroneous

We analyzed subsets of an empirical data set of *Heliconius* butterfly genomes [4] using the same methods as for our simulation study. The log-likelihood curve inferred using the summary method `InferNetwork_ML` for the species *Heliconius melpomene*, *H. hecalesia* and subspecies *H. erato demophoon* was almost horizontal without a noticable shoulder. Given the robustness of summary methods at predicting reticulation number, this result suggests that the the true number of reticulation for this subset is zero (Fig. 9).

**Fig. 9:**
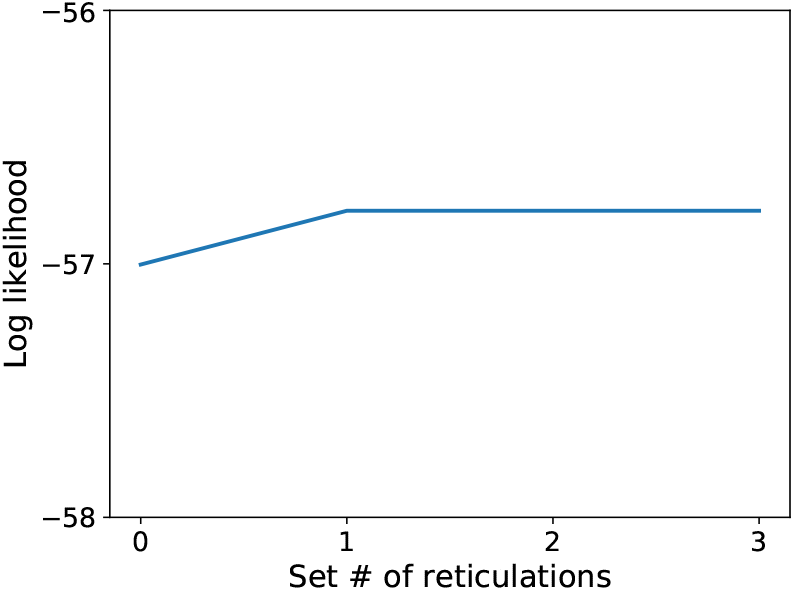
Log-likelihood increase of *Heliconius* species networks identified using the maximum likelihood summary method `InferNetworks_ML`. Gene trees used as input were inferred using `IQ-TREE` from sequence data extracted from an whole genome alignment of *Heliconius* species, pruned to *H. hecalesia*, *H. erato demophoon* and *H. melpomene*.

When using the full Bayesian method `MCMC_SEQ` with the DEO implementation of the DR model enabled, the *maximum a posteriori* phylogeny was a tree without reticulations and an identical topology to the maximum likelihood method with the number of reticulations was set to zero (Fig. 10A,11A). However, when the SR model was used for inference, the full Bayesian method inferred gene flow after speciation from the ancestor of *H. hecalesia* and subspecies *H. erato demophoon* into the *H. melpomene* lineage (Fig. 10B). This was different from the reticulation inferred by `InferNetwork_ML` when one reticulation was allowed, although as mentioned reticulations are unsupported by the log-likelihood curve (Fig. 11B).

**Fig. 10:**
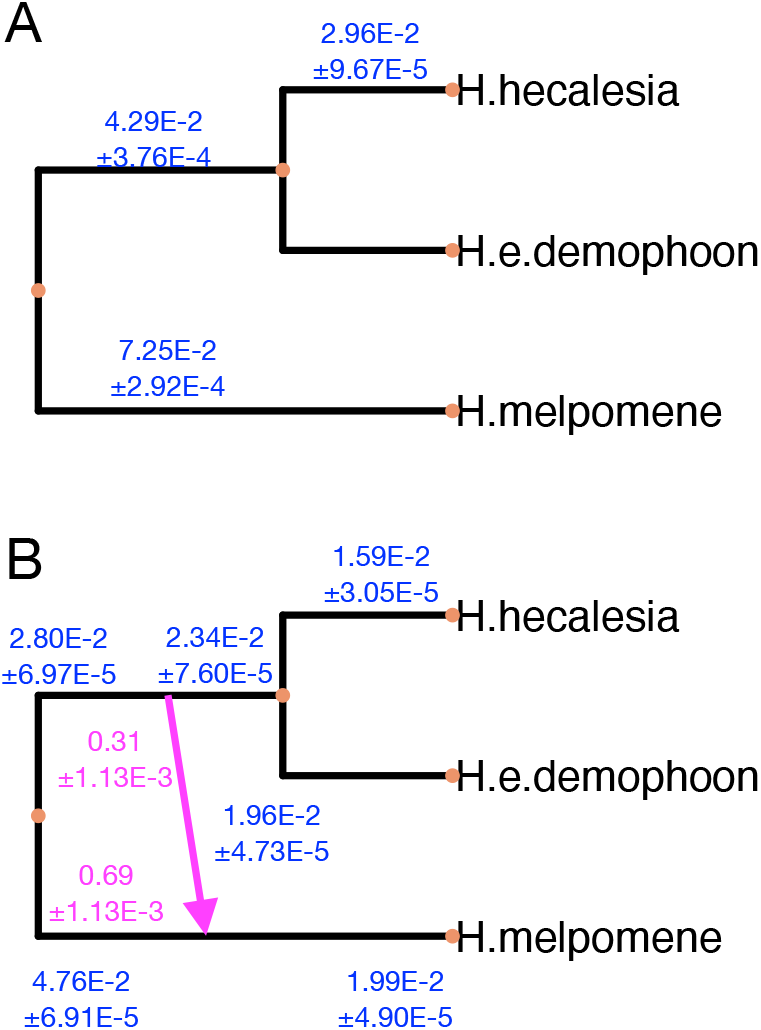
*Heliconius* species networks inferred using the full Bayesian method `MCMC_SEQ`. Phylogenetic inference was performed using the Dirichlet rates model of per-locus rate variation (A) or a single rate model (B), from sequence data extracted from an whole genome alignment of *Heliconius* species, pruned to *H. hecalesia*, *H. erato demophoon* and *H. melpomene*. The posterior expectation and standard deviation of branch lengths are given in expected substitutions per site (blue). The posterior expectation and standard deviation of inheritance probabilities are also shown (pink).

**Fig. 11:**
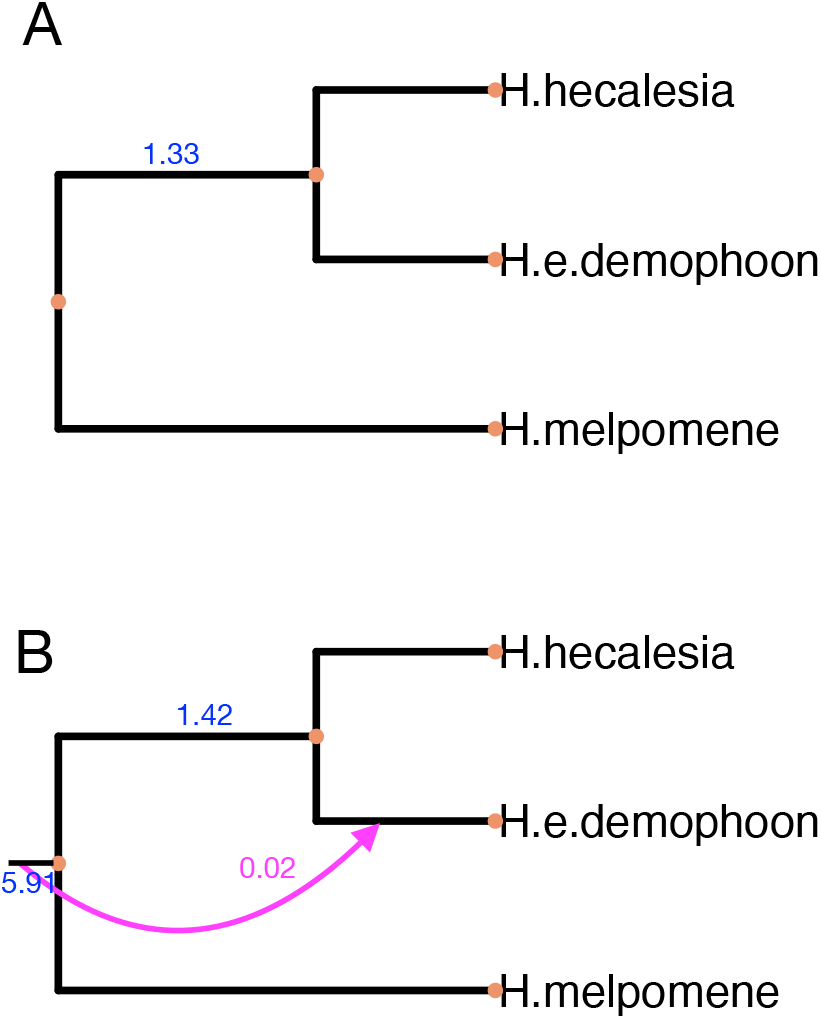
*Heliconius* species networks identified using the maximum likelihood summary method `InferNetworks_ML`. The maximum number of reticulations was set to zero (A), or up to 3 (B). Gene trees used as input were inferred using `IQ-TREE`from sequence data extracted from an whole genome alignment of *Heliconius* species, pruned to *H. hecalesia*, *H. erato demophoon* and *H. melpomene*. Estimated branch lengths are given in coalescent units (blue). Inheritance probabilities are also shown (pink).

Substantial variation in substitution rates was inferred when using the DR model (Fig. S2). Given this observation, the trend we observed in the simulation study where using the SR model leads to the inference of spurious reticulations when rate heterogeneity is present, we suggest this apparent gene flow is an artifact of model misspecification. That is, the true model is one with rate heterogeneity, which is not permitted by the SR model.

The inference of spurious reticulations does not always manifest. We analyzed another subset of three species *H. timareta*, *H. melpomene* and *H.numata*, and no reticulations were inferred by the full Bayesian method regardless of whether the SR or DR models were used for inference (Fig. 12). This is in spite of variation in substitution rates also being inferred for this subset (Fig. S7), but in agreement with the log-likelihood curve and topology inferred using the summary method (Fig. S5A, S6).

**Fig. 12:**
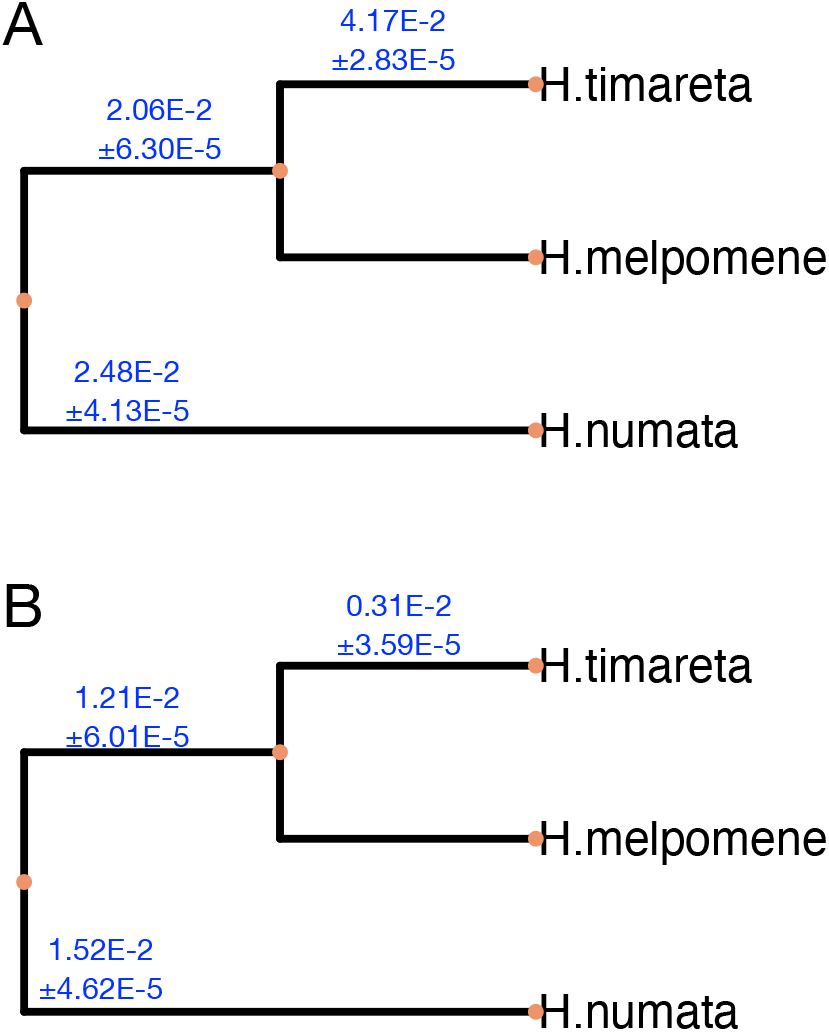
*Heliconius* species networks inferred using the full Bayesian method `MCMC_SEQ` for an alternative subset of taxa. Phylogenetic inference was performed using the Dirichlet rates model of per-locus rate variation (A) or a single rate model (B), from sequence data extracted from an whole genome alignment of *Heliconius* species, pruned to *H. timareta*, *H. melpomene* and *H. numata*. The posterior expectation and standard deviation of branch lengths are given in expected substitutions per site (blue).

Convergence of `MCMC_SEQ` was slower when the DR model was used. For the same number of iterations, the posterior density effective sample size (ESS) was 436 or 1714 when DEO was enabled for the first and second subsets (Fig. S3, S8), compared with 11737 or 2920 when DEO was disabled for the same subsets (Fig. S4, S9).

## Conclusions

Model misspecification is a persistent issue in phylogenetic inference. A major area of misspecification is where gene flow occurs after speciation, which is not accounted for by simple species *tree* models. The authors of this paper, and other developers of methods for systematics, have made progress in relaxing this assumption so that inferred species phylogenies can incorporate gene flow through a species *network* model. However, we have shown that in resolving one area of misspecification, we have made the inference of species phylogenies more susceptible to misspecified substitution models.

Through our simulation study, it is clear that full Bayesian MSNC methods can infer spurious reticulations when the true generative process is one with variation in substitution rates between loci that sequence alignments are derived from, but the model used for inference assumes that a single substitution rate applies to all loci. The accurate inference of gene trees is also harmed by this misspecification, particularly the inference of their branch lengths. For researchers interested in the patterns of substitution rate variation, we have also shown that these rates may to an extent be inferred using the DR model, even in the presence of reticulate evolution. On the other hand, when the true generative process does not incorporate rate variation, using a model for inference that allows for rate variation does not have a substantial negative impact on accuracy. Because of this asymmetry in outcomes, we recommend that species networks should be inferred using the DR or similar model whenever substitution rate variation is possible.

Through our empirical study, we have shown that this misspecification has likely caused spurious inferences when applied to real taxonomic systems. The inference of spurious reticulations may led to incorrect conclusions concerning patterns of reticulation in evolution. This further supports our recommendation that rate variation should be accounted for whenever it may be present. To facilitate our recommendation, we have implemented the DR model in `MCMC_SEQ`, one of the most popular methods for species network inference.

